# A hybrid capture RNA bait set for resolving genetic and evolutionary relationships in angiosperms from deep phylogeny to intraspecific lineage hybridization

**DOI:** 10.1101/2021.09.06.456727

**Authors:** Michelle Waycott, Kor-jent van Dijk, Ed Biffin

## Abstract

Novel multi-gene targeted capture probes have been developed with the objective of obtaining multi-locus high quality sequence reads across any angiosperm lineage. Using existing genomic and transcriptomic data, two independent single assay probe/bait sets have been developed, the first targeting conserved exons from 20 low copy nuclear genes (OzBaits_NR V1.0) and the second, 19 plastid gene regions (OZBaits_CP V1.0). These ‘universal’ bait sets can efficiently generate DNA sequence data that are suitable for systematics and evolutionary studies of flowering plants. The bait sets can be ordered as Daicel-Arbor Sciences custom myBaits. We demonstrate the utility of the bait set in consistently recovering the targeted genomic regions across an evolutionarily broad range of angiosperm taxa.

## Introduction

Target enrichment approaches (eg. Cronn et al., 2012;Lemmon et al., 2012;Weitemier et al., 2014) have developed as a method of choice for systematics and evolutionary studies. Target enrichment probe sets can be designed to efficiently target a large number of phylogenetically informative loci and can be readily scaled up to include the dense sampling that is typical of molecular evolutionary studies. In addition, the method appears to work well with degraded samples, and can be used to obtain data from museum specimens (e.g. Hart et al., 2016) and from environmental samples (e.g. Lentz et al., 2021). While probe sets can be designed to target a narrow set of lineages (e.g. angiosperm families) ‘universal’ probe sets are attractive because of reduced cost and effort as well as the potential to generate broadly comparable data across evolutionarily distant lineages (e.g. Johnson et al., 2018;Breinholt et al., 2021).

In this methods manuscript we present two myBaits target enrichment probe sets that target, 1; conserved exons from 20 low copy nuclear genes and 2; 19 chloroplast gene regions. Leveraging existing genomic and transcriptomic data, these bait sets have been developed with the objective to obtain high quality sequence reads across all angiosperms. These new probe sets can be used to generate DNA sequence data that is suitable for a wide range of systematics and evolutionary analyses for any lineage of flowering plants. The bait sets were tested on a small selection of samples that cover a broad phylogeographic range.

## Methods

### Bait Design and Synthesis OzBaits V1.0

#### Nuclear genes

Target genes were selected from Duarte et al. (2010) who report a set of putatively orthologous low copy nuclear genes shared in *Arabidopsis, Populus, Vitis* and *Oryza* (APVO SSC genes sensu Duarte et al., 2010). For each selected gene (Table 1), we downloaded all coding sequences (CDS) for the corresponding gene family available on Phytozome (*v. 12;* https://phytozome.jgi.doe.gov). We used the CDS for *Arabidopsis thaliana* to retrieve putatively homologous transcript sequences from The 1000 Plants Project (1KP; https://www.onekp.com) using the China National Genebank (https://db.cngb.org) BLAST portal and the following settings: Discontiguous Mega-Blast, expect value=10, maximum target sequences=1000, selected organisms=Magnoliophyta (taxid:3398). The sequences retrieved from the BLAST search were downloaded and combined with the Phytozome data for each gene and made into a BLAST database in Geneious (Kearse et al., 2012; https://www.geneious.com). We queried each BLAST database using the *A. thaliana* gene family member with exon annotations manually added, and the following settings: Discontiguous Mega-Blast, expect value=10, maximum target sequences=1200, results=Hit Table, retrieve=Matching Region with Annotation. We then extracted all sequences matching one or more exon annotations in *A. thaliana* with the caveat that the exon was > 180 bp in length to allow for bait tiling. The extracted sequences were clustered using CD-HIT-EST (Li and Godzik, 2006; http://weizhong-lab.ucsd.edu/cdhit_suite;Huang et al., 2010) with a sequence identity cut-off fraction of 0.95 and a length similarity fraction of 0.2, and one representative sequence (the longest) per cluster was selected. A total of 9,900 representative sequences were used for bait design with 120-nucleotide baits and ∼2X flexible tiling density for a total of 47,001 baits (details of design can be found in Supplemental 1). Bait design (myBaits) and synthesis was performed by Daicel-Arbor Biosciences (formerly Mycroarray; Ann Arbor, Michigan, USA) and has the Design ID **D10121VasPl**.

**Table 1:**
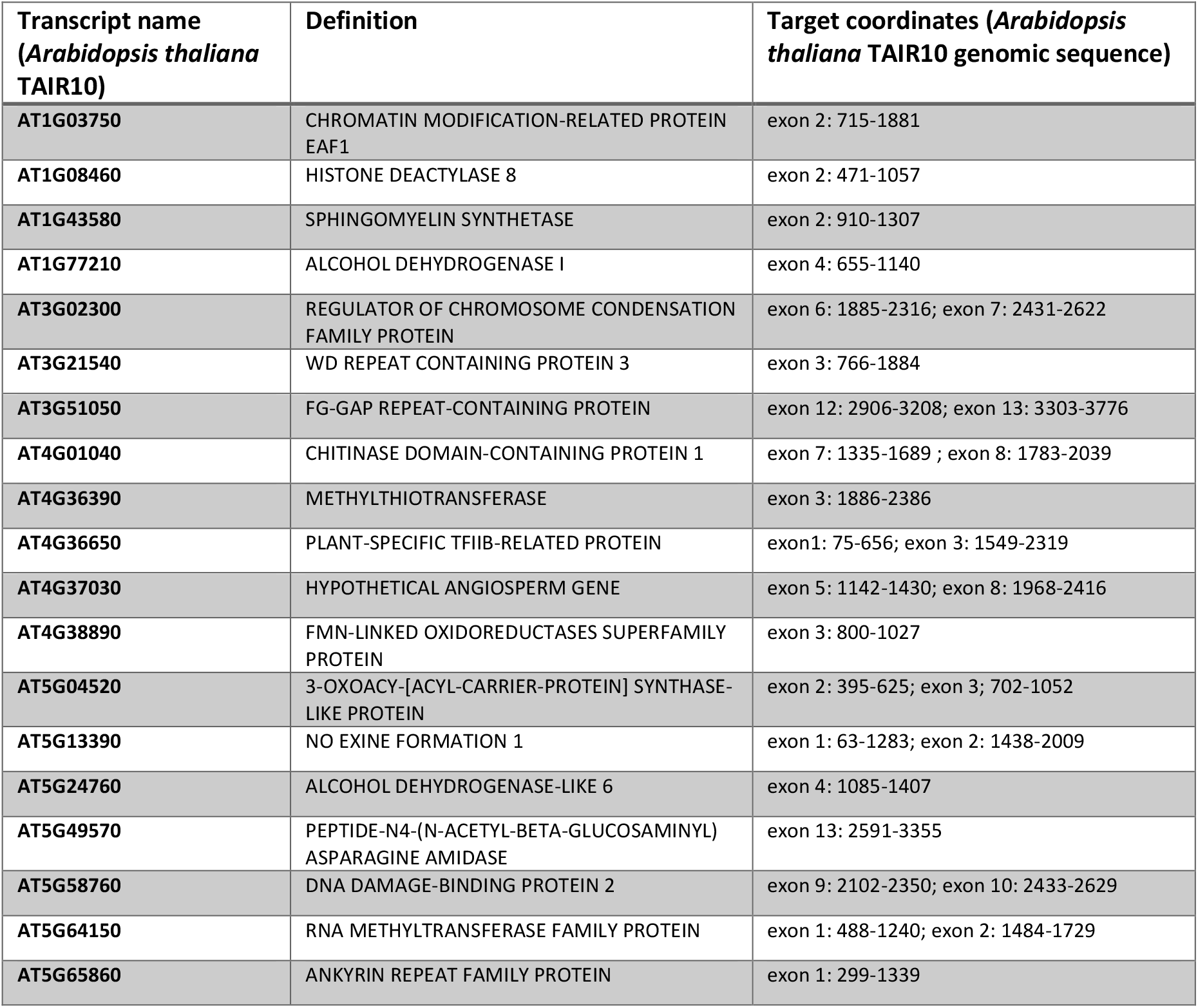
Nuclear low copy genes used for OzBaits_NR V1.0, Design ID **D10121VasPl**

#### Chloroplast genes

Although hybrid capture approaches will frequently recover complete or near complete plastid genomes from off-target sequencing reads, the efficiency of recovery is reduced in degraded material such as herbarium specimens (e.g. Bakker et al., 2016). To improve the consistency of recovery of high-coverage plastid data, we designed a set of probes to target a range of chloroplast regions in angiosperms using the NCBI RefSeq plastid sequences (https://www.ncbi.nlm.nih.gov/genome/organelle/) for probe design. For each target gene, the corresponding CDS were extracted from the RefSeq genome data and clustered as above, using CD-HIT-EST. A total of 2978 sequences, targeting 19 plastid regions (Table 2) were used for bait design with 120-nucleotide baits and ∼2X flexible tiling density for a total of 14,922 baits (details of design can be found in Supplemental 2). Bait design (myBaits) and synthesis was performed by Arbor Biosciences (formerly Mycroarray; Ann Arbor, Michigan, USA) and has the Design ID **D10122Plstd**.

**Table 2:**
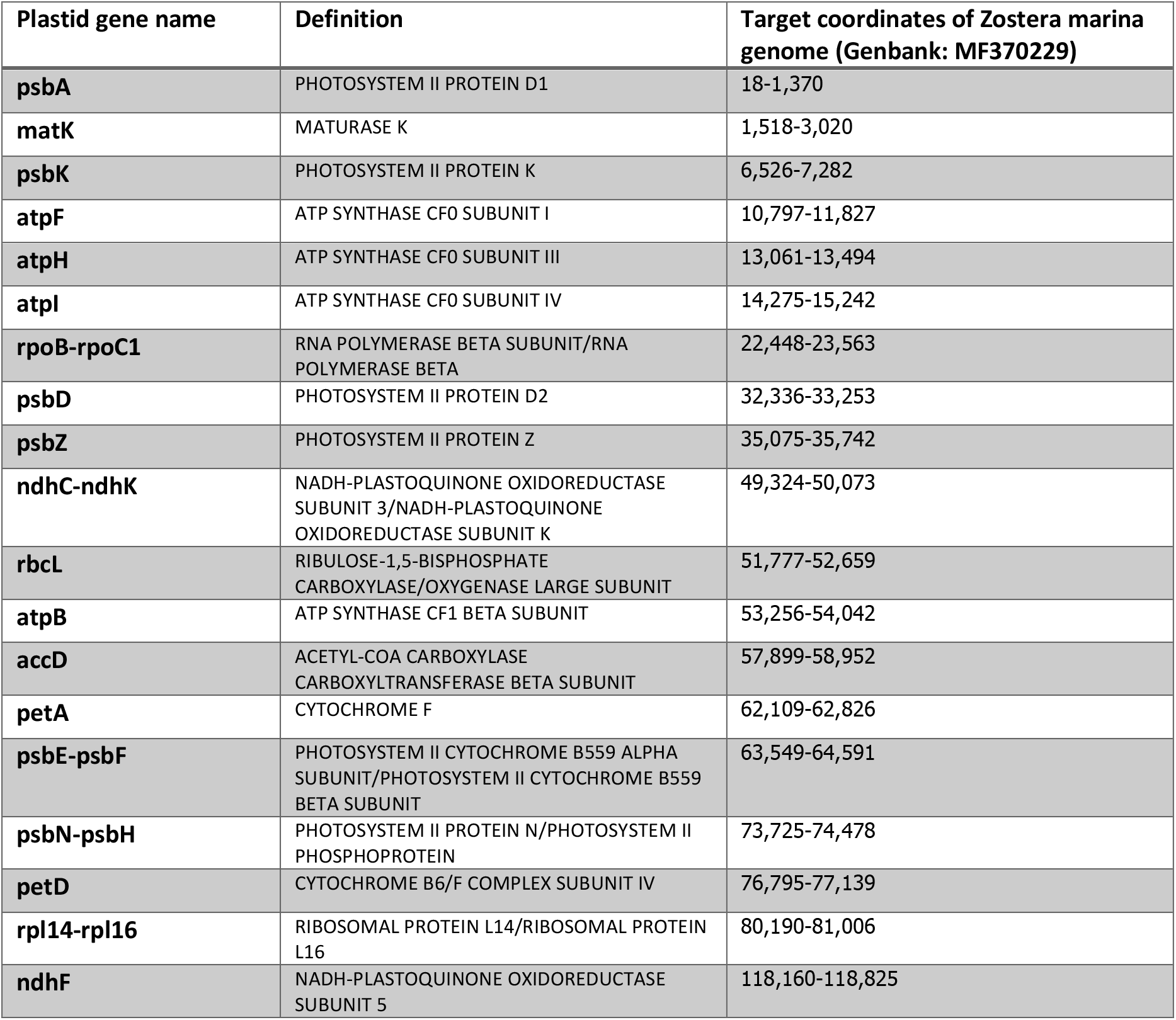
Chloroplast genes used for OzBaits_CP V1.0, Design ID **D10122Plstd**

### Library Preparation and Capture

DNA for targeted capture needs to be of the best quality possible, but this approach has also been successfully tested on degraded DNA (herbarium material from the late 19^th^ century) although sequencing success varies. Most of the libraries prepared in our laboratory used DNA that was extracted with the Qiagen Plant Mini kit. DNA is then normalized to 2ng/uL before proceeding to library preparation, DNAs with lower concentration were used neat, but sequencing success generally decreased. A detailed description of laboratory procedures can be found in Supplemental 3. The described method follows the NEBNext^®^ Ultra™ II DNA Library Prep Kit (v4.0) with Sample Purification Beads Kit (New England Biolabs, Ipswich, Massachusetts, USA). But other kits can be used too, adjustments need to be made to the protocol though. To maintain consistency in the library preparation all liquid handling was done on an Eppendorf epMotion^®^ 5075t - Liquid Handling Workstation, but steps can also be performed manually, both approaches are described in Supplemental 3. To keep cost down and to be able to multiplex in high sample numbers custom stubby Y-adapters were developed and used in the library perps described in Supplemental 3. A detailed description of development with necessary sequence files can be found in Supplemental 4 - 7.

After library preparation, samples are pooled to reduce costs of Hybrid Capture, in Supplemental 3 pools are made with 16 libraries (step 44-47), but if samples are of low quality pools in sets of 8 will generally increase success, or, if you are dealing with ancient DNA you might consider doing captures on single libraries and follow ancient DNA workflows.

Hybrid Capture was performed following the manual provided by myBaits. Our captures were performed at 65°C, but temperatures can be changed depending on the capture efficiency needed. During the hybridisation step Chill-out™ red liquid wax was added to avoid evaporation. After capture libraries were amplified with indexed primers (Supplemental 6), cleaned up with Ampure and pooled to equimolar concentrations (Nuclear and Chloroplast separately).

Chloroplast DNA captures are generally orders of magnitude stronger in molarity. This is probably due to the higher DNA extraction yield (number of genome copies recovered) and fewer loci that are used for capture. Only 10% (molarity) of the final library pool was allocated to the Chloroplast library. As a final quality step libraries are size selected according to insert sizes and sequencing platform used to avoid dimers and over-long inserts. If DNA is expected to be degraded size selection should target smaller inserts but avoid primer dimers. In our laboratory we generally sequence on an Illumina NovaSeq S1 flow cell pooling 3-5 plates worth of samples.

To test the bait sets OzBaits_NR V1.0 and OzBaits_CP V1.0 a small number of samples covering a broad range of angiosperms was tested (Table 3, Supplemental Table S8). Plant material was collected form either dry herbarium specimens or from the Adelaide Botanic Gardens living collection. Approximately 20-40 mg of dry material was transferred to a 2mL screw cap tube and pulverized in an Omni Bead Ruptor 24. Total genomic DNA was extracted using the DNeasy™ Plant Kit (Qiagen) according to the manufacturer’s instructions. Library prep was performed according to Supplemental 3 and targeted capture at a hybridization temperature of 65°C.

**Table 3:**
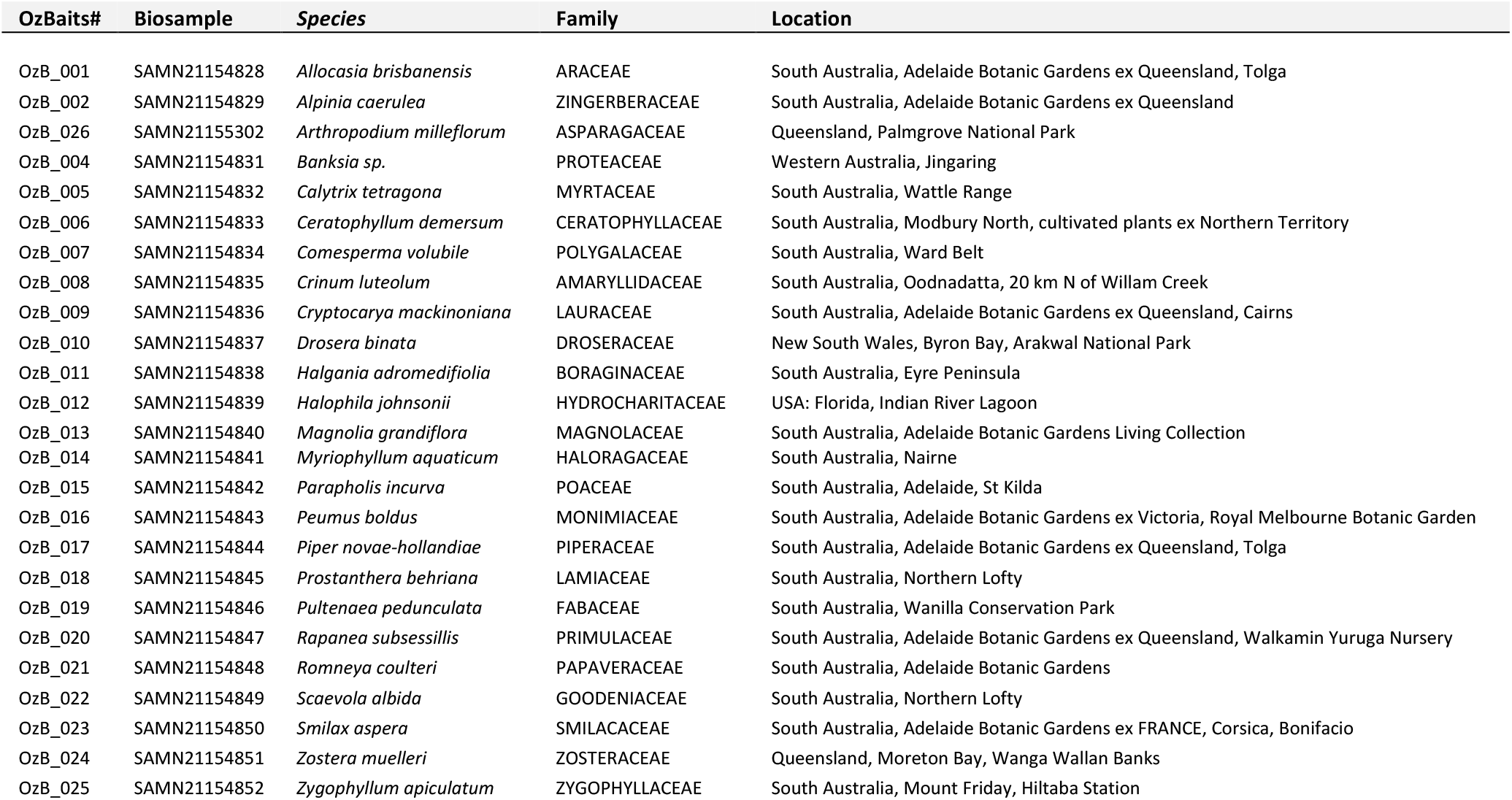
Samples used to test OzBaits_NR and OzBaits_CP V1.0. Detailed information can be found in supplemental Table S8. BioSamples are part of NCBI Bioproject PRJNA680375.

### Data processing and analyses

High-throughput 150 bp paired-end reads were processed using CLC Genomics Workbench (v. 7; **https://www.qiagenbioinformatics.com/**). De-multiplexing, quality trimming (using a Phred score threshold of 20) and de-novo assembly (length fraction=0.7, similarity fraction=0.9, minimum contig length=300) was performed to obtain contigs at each locus (low coverage cut-off = 10, insert Ns for regions of low coverage, conflict resolution = vote). To identify the ‘on target’ nuclear contigs from each assembly we first made a BLAST database comprising the full set of reference genes from the Phytozome dataset, and we queried against these using the contigs from each *de-novo* assembly. Contigs corresponding to target loci were then retrieved by creating a BLAST database for each *de-novo* assembly and running a query against the best matching Phytozome reference sequences identified in the first step, using an E-value ≤1E-100 and maximum target sequences set to 20. For each sample, including a diverse set of angiosperm taxa (Table 3, Supplemental Table S8), we assessed per locus recovery, gene copy number (assuming that overlapping and non-identical contigs for a given gene represent paralogues) and the total length of sequence recovered per sample (Table 4).

**Table 4:**
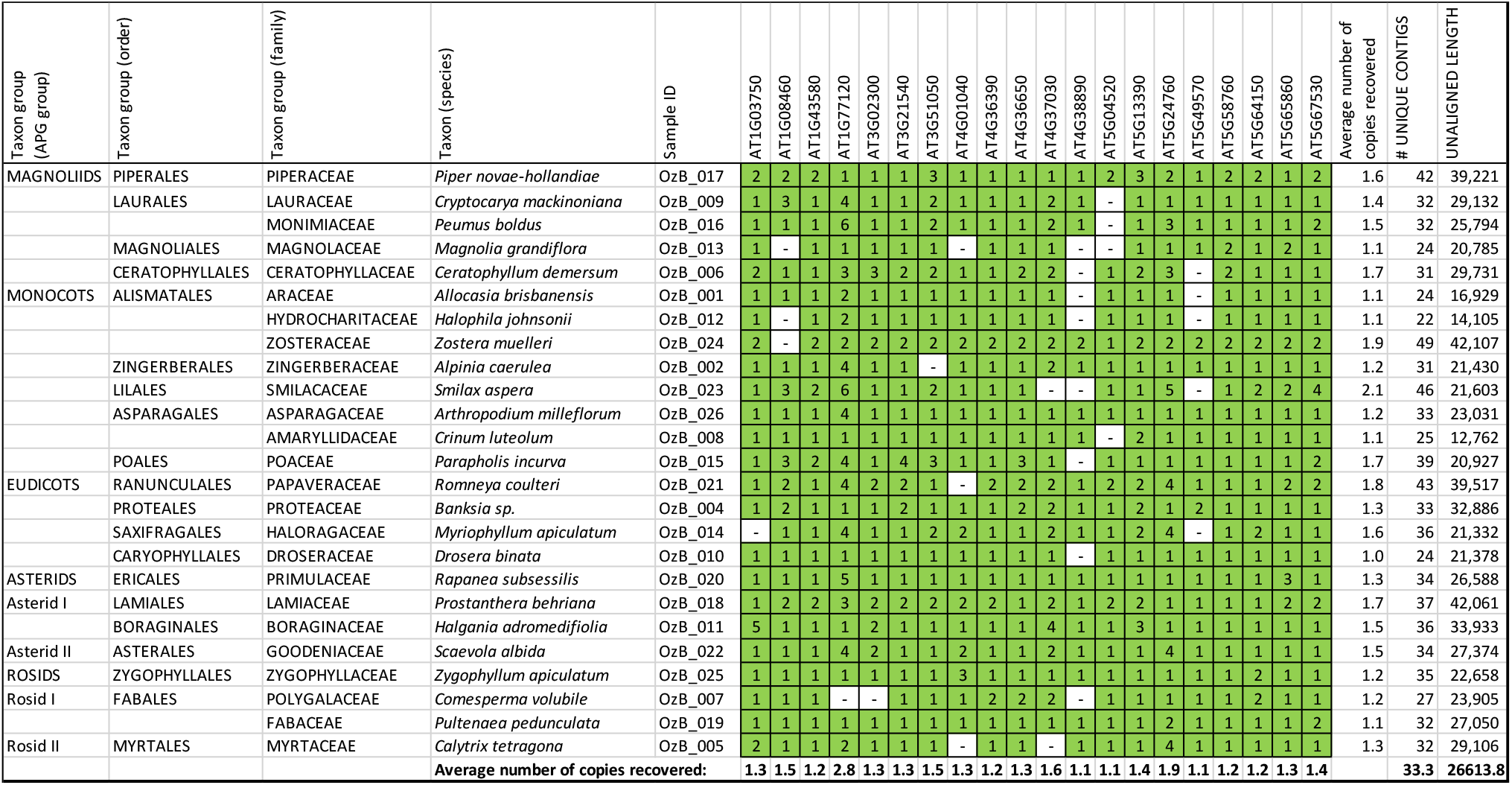
Recovery success of tested samples on low copy genes in bait set OzBaits_NR v1.0. The average recovery of genes was 0.942 with an average of 33.32 unique contigs and an average length of 26,614bp.

In order to recover the targeted plastid regions, we used the chloroplast genome sequence of *Arabidopsis lyrata* (KX886355) as a BLAST reference to query the sample specific *de novo* assemblies (as above) using an E-value ≤1E-20 and maximum target sequences set to 20.

## Results and Discussion

Recovery of target loci across a diverse set of angiosperm samples was high for both the nuclear (Table 4) and chloroplast target regions (Table 5). For the targeted nuclear gene regions, there appears to be little evidence of a taxonomic effect on gene recovery, suggesting that these should be effectively recovered for any angiosperm. Plastid gene regions are rarely targeted given that these are commonly recovered from off-target reads (e.g. Weitemier et al., 2014). However, a plastid specific probe set enables consistent recovery of the target regions from a wide range of samples such as environmental and ancient DNA (e.g. Schulte et al., 2021) comprising degraded samples with typically low recovery of genomic DNA. Both chloroplast and nuclear gene regions have been successfully employed in evolutionary analyses at a range of taxonomic levels (Nge et al., 2021;Waycott et al., 2021).

**Table 5:**
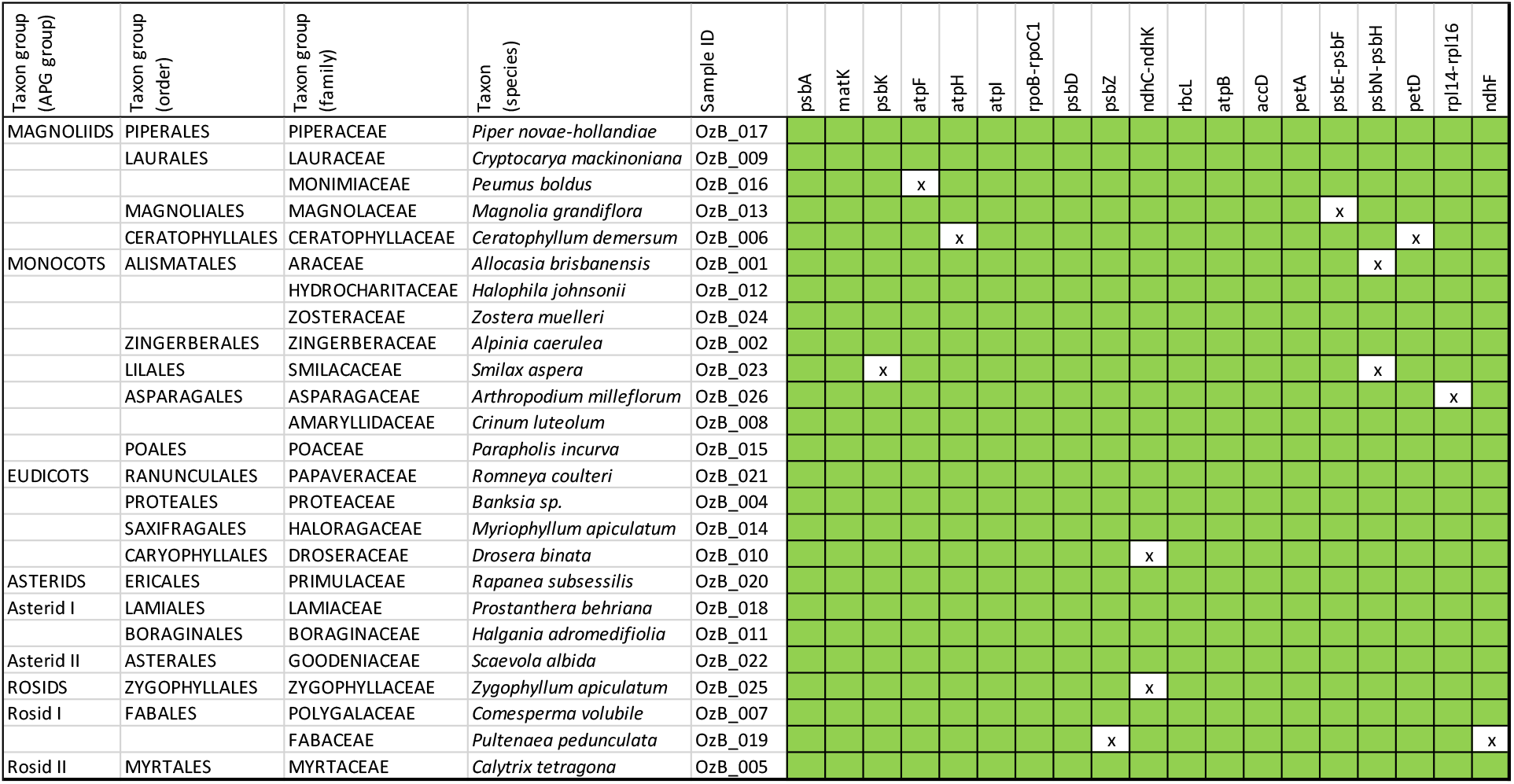
Recovery success of tested samples on plastid genes in bait set OzBaits_CP v1.0. For many of the samples an almost complete genome was recovered.

