## Supplemental 3 for "A hybrid capture RNA bait set for resolving genetic and evolutionary relationships in angiosperms from deep phylogeny to intraspecific lineage hybridization"

**PROTOCOL**
**Library perpetration for Hybrid Capture**

The goal of this procedure is to prepare fragmented DNA libraries having unique Illumina TruSeq adapters. These libraries will subsequently be enriched using solution hybridization, also known as Target Enrichment or Hybrid Capture (e.g. see Gnirke *et al.* 2009 and Blumenstiel *et al.* 2010) using myBaits® RNA probes. This protocol has been written to be performed without an automatic liquid handler. However, the pre-capture library preparation for this study was done on an Eppendorf epMotion® 5075t - Liquid Handling Workstation with 50ul and 300ul single and 8 channel tools with filter tips and a gripper. Details for robot liquid handling are described in boxes. The final construct of library with insert after capture is shown in Figure S1, additional details can be found in Supplemental ##.

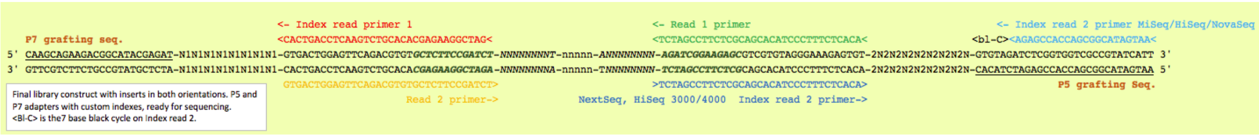

*Figure S3.1 Library construct after Hybrid Capture with Illumina sequencer adapters see Supplemental 2 for details.*

**Original sources**
Library preparation NEBNext® Ultra™ II DNA Library Prep Kit (v4.0) with Sample Purification Beads Kit (New England Biolabs, Ipswich, Massachusetts, USA) and Hybrid Capture protocol with MyBait RNA Hybrid Capture Kit v.4 (Arbor Biosiences, Ann Arbor, Michigan, USA)

**Materials**

Essential

Library Preparation (– equipment used in our lab)
- Sonicator for shearing samples – Bioruptor® Pico
- Pipettes single channel P10, P20, P200, P1000; multichannel P20 and P200
- Vortex mixer
- Mini centrifuge with adapters for 1.5 mL and 0.2 mL tubes/strips
- Rare-earth 96-well magnet stand (Pre-PCR) – Alpaqua FLX 96 well
- Rare-earth 1.5mL tube magnet stand
- Thermal cycler – Eppendorf™ Mastercycler™ Nexus Thermal Cycler
- Agarose gel system – Biorad Wide Mini-Sub Cell GT Cell
- Gel Documentation System – Axygen GD-1000
- Automated electrophoresis system for DNA quality control – Agilent 4200 TapeStation System with Agilent High Sensitivity D1000 ScreenTape

- Sonicator vials – Bioruptor 600µl tubes
- 96-well PCR plates – Eppendorf twin.tec® PCR Plates LoBind
- Plate Sealing Film for PCR – Biorad Microseal® 'B' PCR Plate Sealing Film
- 1.5 mL low bind tubes

- Plate Sealing Film – Axygen® PCR-AS-600 PCR 50µm Aluminium Sealing Film

- AMPure XP beads or substitute (NEBNext Beads)
- Molecular grade Ethanol 100%
- 1X TE (10 mM Tris pH 8.0, 1 mM EDTA)
- 5 M NaCl
- 1 mM Tris-HCl (pH 8.0), 100 µM EDTA, 50 mM NaCl
- Agarose molecular grade
- 1X TBE
- Library prep Kit – NEBNext® UltraTM II DNA Library Prep with Sample Purification Beads Library Preparation Kit
- Proof reading polymerase – NEBNext® Ultra™ II Q5® Master Mix
- Illumina TruSeq-style stubby Y adapters with 8nt identification Tags (for custom design see Supplemental Table S3)
- Loading dye (6X) and size standard for agarose gels
- Amplification primers (@ 100µM) for Illumina adapters (includes phosphorothioate bonds (*))
  P7_preCap_Long 5’ ACACTCTTTCCCTACACGACGCTCTTCCGATC*T 3’ (65.5°C melt)
  P5_preCap_Long 5’ GTGACTGGAGTTCAGACGTGTGCTCTTCCGATC*T 3’ (65.8°C melt)

Hybrid Capture (additional items)

- Rare-earth 96-well magnet stand (post PCR)
- Water bath or incubation oven capable of 65°C
- Heat block capable of 65°C

- 8-well strip cap tubes with attached lids

- Custom blocker oligonucleotides (1000 µM) to replace myBaits Block A (Supplemental Table S4)
- PCR primers for amplifying your sequencing libraries after capture (includes 3’ phosphorothioate bonds (*)). Indexed primers P5 and P7 (@ 100µM) for finishing Illumina adapters (includes phosphorothioate bonds (*)) for custom design see Supplemental Table S4
  P5_primer_Idx### 5’ AATGATACGGCGACCACCGAGATCTACAC-NNNNNNNN-ACACTCTTTCCCTACACGA*C 3’
  P7_primer_Idx### 5’ CAAGCAGAAGACGGCATACGAGAT-NNNNNNNN-GTGACTGGAGTTCAGACGTG*T 3’
- 10 mM Tris-Cl, 0.05% TWEEN®-20 solution (pH 8.0-8.5) (30 µL per enrichment reaction)
- KAPA HiFi HotStart Ready Mix PCR Kit
- Chill-out™ red liquid wax

Useful

- Multichannel pipettes (P20, P200)
- Automatic liquid handler – Eppendorf epMotion® 5075t - Liquid Handling Workstation with 50ul and 300ul single and 8 channel tools with filter tips
- Sage Science Pippin Prep with 2% agarose DNA gel cassettes (100bp-600bp) or 1.5% agarose DNA gel cassettes (250bp-1.5kb)

### Adapters

If you are using custom sequencing adapters (i.e., not the standard Illumina TruSeq adapters) and something like adapters described in Supplemental 4, and you have ordered both P5 and P7 oligos from your preferred oligonucleotide supplier, you need to prepare (anneal) the tagged adapters. This is done by mixing the P5 tag### with the complement P7 tag###, and heating these up and letting them cool down slowly to make double-stranded stubby Y-adapters. This needs to be done for all Y-adapters and it is best/easiest done in a PCR plate with a thermal cycler.

1. Assemble the following components to make Annealing Buffer:

   9.9 mL 1X TE (10 mM Tris pH 8.0, 1 mM EDTA)
   100 µL 5 M NaCl
   ______________________________

   10 mL Total

2. Using the Annealing Buffer, assemble the following components to create the adapters:

   25 µL Annealing Buffer
   12.5 uL 100 µM P5 Adapter tag###
   12.5 uL 100 µM P7 Adapter tag###
   ______________________________

   50 uL Total 25µM Adapter

3. Anneal:

   95°C for 1 minute
   ⇨ -0.1°C /sec for 800 seconds (13.33 minutes)
   ⇨ 14°C hold

4. Seal plate with aluminium seal and store adapters at -20°C

### Normalization and Fragmentation

5. Equilibrate DNA samples to ~ 2.0 ng/µL in a volume of 100µL (if possible, do this on liquid handler (e.g. epMotion®)) in a 96-well plate
6. Transfer DNA from plate to 600µl Bioruptor tubes and keep plate.

Normalization for Eppendorf epMotion® 5075t was setup with Normalization Calculator provided by Eppendorf. CSV file with individual pipetting volumes were imported into robot.

Shear DNA to a size distribution peaking around 300-500 bp using sonicator (for this study a Diagenode Bioruptor Pico was used).

Our Diagenode Bioruptor Pico run cycle was 15 Sec On, 90 Sec Off and 6 repeats

7. Return sonicated DNA to the 96-well plate seal with aluminium foil and store sonicated DNA at -20°C for future use

*The most important pre-requisite for any NGS library preparation is high-quality DNA. Sample handling and DNA isolation procedures are therefore critical to the success of the experiment. Residual traces of proteins, salts or other contaminants could degrade the DNA or decrease the efficiency of the enzymatic activities necessary for optimal library preparation.*

*It is not essential to have a sonicator, you can also do the library preparation with the Fragmentase® enzyme (NEB). We have done trials with the NEBNext® Ultra™ II FS DNA Library Prep Kit (1/3 volume) and had very good results. Fragmentase incubations of 5 min seem long enough for good Herbarium DNA samples.*

#### ***NEBNext Ultra II DNA Library Preparation***

This library preparation is an adaptation of the NEBnext Ultra II DNA Library Prep manual as the End Prep and Ligation reactions were performed in 1/3d of the recommended reaction volumes. Other library preparation kits can be used too, but steps need to be adapted according to supplier's manual.

#### End-Prep

*This step repairs the ends of the DNA after sonication as there may be fragments that are partly broken. This step also phosphorylates DNA ends. If you are using NEBNext® Ultra™ II FS DNA Library Prep Kit the fragmentation will occur in this step too.*

For runs on epMotion® 5075t Eppendorf LoBind twin.tec® PCR Plates were used so gripper could move the plates on the deck. Due to the dead volume in 1.5mL tubes an excess of master mix is required; 5 extra reactions were added. Automated pipetting was done at very slow speeds due to the viscosity of buffers. Tips were rinsed in DNA solution at bottom of plate by mixing 5x. Both master mix and plate with fragmented DNA were kept at 4°C on robot's thermoblocks. Once program finished plates were placed in thermal cycler by hand.

8. Remove the NEBNext Ultra II End Prep Enzyme Mix reagents (Green cap) from storage (-20 °C) and allow them to thaw on ice. Take plate with fragmented DNA out of freezer and let thaw. Spin at max speed for 1 minute before use
9. Pipette 16.7 µL of fragmented DNA into new low bind PCR plate with multichannel P20 pipette and place plate on ice
10. Before use spin liquids down and pipette the entire volume up and down at least 10 times to mix thoroughly and then spin again
11. Prepare reaction on ice using the volumes shown in Table 1 in a single 1.5mL tube and mix by pipetting up and down. Spin and keep on ice

*Table S3.1 End-Prep reaction*

| *Cap Colour* | *Reagent* | *As per manual* | *1X Master Mix (1/3 of original NEBNext II Kit)* | *#samples+1 Master Mix* |
|---|---|---|---|---|
| | Fragmented DNA | 50 µl | 16.7 µL | - |
| Green | NEBNext Ultra II End Prep Reaction Buffer | 7.0µl | 2.33 µL | |
| Green | NEBNext Ultra II End Prep Enzyme Mix | 3.0 µl | 1.0 µL | |
| | | 60 µL | 20 µL | **Dispense 3.3 µl** |

*Caution: The End-repair buffer is very viscous. Care should be taken to ensure adequate mixing of the reaction.*

12. Dispense with P10 pipette 3.3 µL of master mix into each well of plate with fragmented DNA, change tips for each sample
13. Seal plate with Biorad Microseal® 'B' spin in centrifuge and place plate in PCR machine
14. Run End Prep program: Incubate for 30 min at 20°C then 30 min at 65°C and cool down to 4°C

    If thermocycler allows it, set the heated lid to 75°C
15. Return plate to ice or thermoblock at 4°C

### Adapter Ligation

*Up until this point DNA has been repaired and an A-tail added to the 3' ends of the DNA fragments, next steps will ligate a stubby Y-adapter (Supplemental Table S5) onto these ends.*

In this study we used a stubby Y adapter stock of 25µM resulting in a final concentration of 1.44 µM (this is ~1:15 volume ratio (25µM Adapter: Total Volume)). If concentrations of DNA are low a 2.5µM adapter solution would reduce dimer formation in subsequent PCR (it is recommended to do some concentration tests to find the right molarity). 96 stubby Y adapters were synthesised (Supplemental 5, trcY_Tag001 - trcY_Tag096) and stored in a plate at -20°C.

16. Remove the ligation reagents (Red cap) from storage (-20 °C) and allow them to thaw on ice. Briefly spin and mix the Ultra II Ligation Master Mix by pipetting up and down several times
17. Prepare adapter-ligation mix by assembling reagents on ice according to Table 2. Ensure optimal mixing by pipetting up and down

> For epMotion® 5075t work in multiples of 8, as multichannel pipetting tool will speed-up the movements. Due to the dead volume in the 1.5mL tubes more master mix is required; 3 extra reactions were added to mater mix. Once program has finished place plate in thermal cycler by hand.

*Table S3.2 Adapter Ligation Reaction Mix*

| ***Cap Colour*** | ***Reagent*** | ***As per manual*** | ***1X Master Mix.*** *(1/3 of original NEBNext II Kit)* | ***#samples+1 Master Mix*** |
|---|---|---|---|---|
| | End-Prep reaction line 15 | 60µL | 20µL | - |
| | NEBNext Ultra II Ligation Master Mix | 30µL | 10µl | |
| | NEBNext Ligation Enhancer | 1µL | 0.33µl | |
| | Adapters (25µM) | 2µl | 2µl | Each sample with unique adapter |
| | Total | 93µl | 32.33 µl | **Dispense 10.3** µl |

18. With P20 pipette dispense 10.3 µl of master mix into End Prep reaction (line 15), change tips for each sample
19. Pipette 2µl of Adapters (25µM) into each well using P20 multichannel pipette
20. Seal plate with new Biorad Microseal® 'B', spin in centrifuge and place plate in PCR machine
21. Incubate at 20°C for 20 min in thermal cycler with heating lid turned off

*NOTE: it is recommend performing the clean-up step immediately after ligation. However, if you intend to stop after ligation without the clean-up, it is suggested to store DNA overnight at -20°C. The clean-up step can be continued on the following day without affecting the quality or the yield of the library.*

### Post-Ligation Clean-up without Size Selection

*This step is crucial to remove un-ligated adapters and fragments with ligated adapters that are too small for sequencing. For Hybrid Capture we are targeting inserts between 250-600bp. Keep in mind that adapters will add ~84bp to this intermediate stage of the library.*

Allow NEBNext Sample Purification Beads (AMPure XP or equivalent) to equilibrate at room temperature for at least 30 min. Mix beads thoroughly (vortex) to ensure homogenous resuspension. Make enough fresh 70% Ethanol. If possible, use multichannel pipettes for clean-up steps.

22. Add 32 µL (1.0X) of homogenous (AMPure XP) beads to each adapter-ligated DNA sample. Mix well by pipetting up and down 10X
23. Incubate the mixture for 5 minutes at room temperature

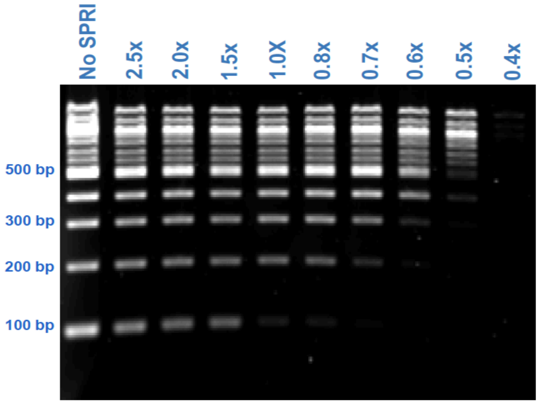


*Figure S3.2 Guideline for AMPure XP selection range according to AMPure XP concentration. (Source: https://www.broadinstitute.org/files/shared/illuminavids/SamplePrepSlides.pdf)*

24. Put the tube(s)/plate on the rare-earth magnet stand to capture the beads. Let sit for about 5 minutes or until solution is clear
25. Using a separate filter-tip for each tube/well, aspirate liquid from tubes and discard
26. Add 150 µL 70% Ethanol to each tube in the stand
27. Incubate 30 seconds and remove the Ethanol
28. Repeat wash steps 26 and 27
29. After wash, remove all residual ethanol without disturbing the beads. Use P20 or P10 pipettes to aspirate small volumes of residual ethanol
30. Air-dry the beads at room temperature for 3-5 min or until the residual ethanol has completely evaporated

> This clean-up protocol with following PCR setup was created as a separate epMotion® program and was run independently. This is one of the most tip consuming parts of the protocol and we recommend the reuse tip function where possible. Before you start make sure you add plenty of 70% ethanol, 0.1 TE and AMPure XP (or other type of PEG-bead clean-up system) to the reservoirs, these have a large dead space. Let all liquids come to room temperature. Make sure AMPure XP is mixed well before you start the program. Pipetting speeds should be very slow to avoid tips picking up beads with DNA. Do not perform a pre-wetting step when pipetting ethanol out of plates, this will resuspend the beads off the magnet. The PCR setup (next section) was part of same liquid handling protocol.

*IMPORTANT: Do not over-dry the beads as this will decrease yield. The bead pellet is dry when the appearance of the surface changes from shiny to matt.*

31. Remove tube(s)/plate from the magnetic stand. Add 18 µL of 0.1xTE to the bead pellet, mix well by pipetting up and down. Incubate for 3 min at room temperature. Spin tube(s)/plate very shortly to collect all liquid and place tube(s)/plate back on magnetic stand for 5 min or until the solution is clear
32. Proceed promptly to preparation of Master Mix for Amplification of Adaptor-ligated DNA
33. Leave plate with cleaned-up products with beads on magnet

*SAFE STOPPING POINT: Adapter-ligated DNA can be stored at -20°C up to 1 week. Remove library off beads first.*

### Amplification of Adaptor-ligated DNA

*In this step the library will be amplified to an intermediate stage (see Supplemental 4, 1st round of PCR Figure S4.1) to ensure it is at a high enough molarity to perform a Hybrid Capture reaction. Do not over amplify libraries as this will increase clonality of reads. The priming sites on the Y-adapters will result in each insert having a partial P5 and a P7 adapter.*

#### Primer preparation

34. Prepare working stock of Primer Mix by diluting the PreCapture_long primers (100 µM) in nuclease-free water to the final concentration of 10 µM each
35. Store at -20°C and thaw on ice before use

*PCR Set-Up (NEB Next Ultra II Q5 MM)*

The following steps are done in a larger volume than the 1/3 reaction volumes used in previous steps. Additional NEB Next Ultra II Q5 MM needs to be purchased as the kit does not provide sufficient 2xMM.

On epMotion® 5075t PCR was setup in to stages of 48 samples. Master Mix was prepared in two 1.5mL tubes of 50 reactions (48+2) each. MM was dispensed in final PCR plate first and cleaned-up library was then dispensed into wells with multichannel pipetting tool.

36. Take PCR reagents from -20°C and allow them to thaw on ice. Briefly vortex and spin down each reagent before use
37. Assemble the following reaction on bench using the volumes shown in Table S3.3. Ensure optimal mixing by pipetting up and down. If doing a 96-well plate worth of samples split Master Mix over 2 x 1.5mL tubes

*Table S3.3 Library amplification reaction*

| ***Cap colour*** | **Reagent** | **Volumes MM 1X** | **X Master Mix +2** |
|---|---|---|---|
| | Purified adapter-ligated library from step 35 (30uL) | 16.5µL library | |
| | NEB Next Ultra ii Q5MM | 17.5 µL | |
| | Primer Mix (10 µM each PreCap long primers) | 1 µL | |
| ***Total*** | | 35 µL | **Dispense 18.5 µl** |

38. Dispense 18.5 µl of master mix to each tube or well in a PCR plate
39. Carefully take 16.5 µl of adapter-ligated library from step 33 (plate with beads still on magnet) and pipette into PCR tube/plate. Use multichannel P20 with filter-tips if possible
40. Seal plate with new Biorad Microseal® 'B', spin in centrifuge and place plate in PCR machine. Perform amplification according to conditions of Table S3.4

*The libraries have now been amplified and are high copy. The following steps will be done by pipetting manually and should happen in a Post PCR laboratory.*

*Table S3.4 Cycling conditions for NEB Next Ultra II Q5MM for Hybrid Capture.*

| **Temperature** | **Time** | **Cycles** |
|---|---|---|
| **98 °C** | 30 sec | 1 |
| **98 °C** | 10 sec | 17 Cycles |
| **65 °C** | 30 sec | |
| **72 °C** | 30 sec | |
| **72 °C** | 2 min | 1 |
| **4 °C** | Hold | |

*SAFE STOPPING POINT: Adapter-ligated amplified DNA can be stored at -20°C until needed.*

*At this stage of the library-prep process an amplified and purified DNA library with internal 8bp barcodes has been created with inserts that are 200bp or larger. The flanking regions of inserts are partial P5 and P7 adapters with no indexes and grafting sites. As these are PCR products it is recommended to continue work in a post PCR lab.*

### Visualization, quantification and pooling of samples

#### Visualization

41. Prepare a 1.5% agarose gel to check amplification of libraries
42. Mix 3 µl of library with water and loading dye. Load gel with libraries and size standards and run for approximately 40 minutes at 80V (voltage and time will depend on gel-size)
43. Visualize on Gel Documentation System and save gel image

If library prep was successful a faint DNA-smear (300bp and larger) should be visible for each sample.

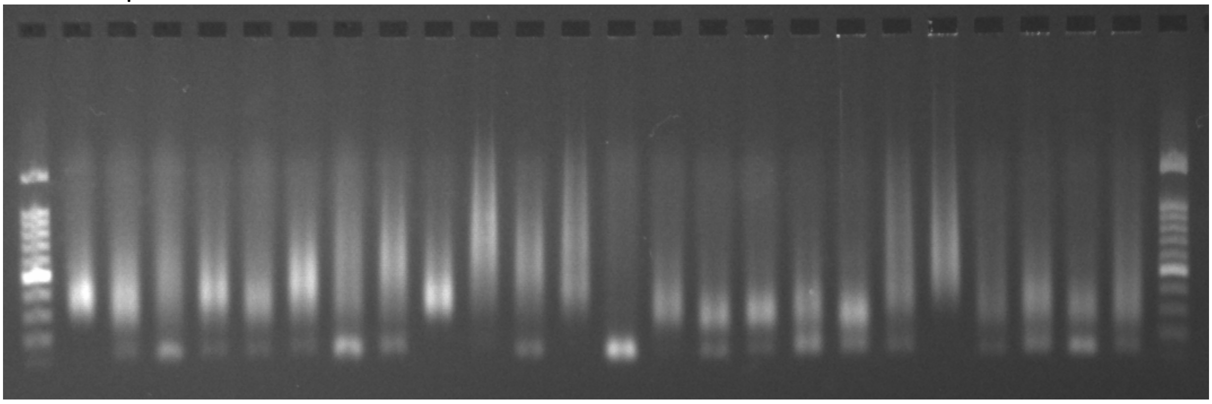

*Figure S3.3 Example of 1.5% agarose gel with amplified libraries.*

*The Hybrid Capture will be performed on pools of libraries. The number of samples in a pool will depend on nature of the project and the number of internal 8bp barcodes (Supplemental Table S5) used in the ligation step (step 19). For this study 16 libraries were pooled for each Hybrid Capture reaction.*

#### Quantification

*There are many ways the individual libraries can be quantified. Automated electrophoresis tools such as the Agilent TapeStation or Perkin Elmer LabChip GX are excellent high precision technologies. In the development of this protocol, we used both these approaches, but a simple visual assessment of the gel seemed to do an equally good job.*

44. Grade each lane of gel with a score of 1 to 4
45. 1 = best/most DNA 4= worst/least DNA in the 300-650bp range
46. The scoring is done relative to the strongest samples in the pool and not to the ladder
47. Pipette libraries of each pool into Eppendorf low bind tube using filter tips

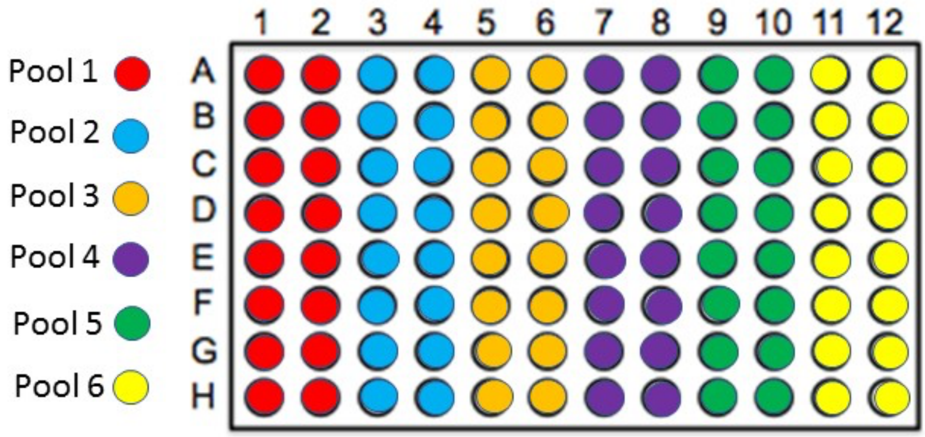


*Figure S3.4 Pooling setup for a 96-well plate of libraries using the scores from step 44. Volumes pipetted: Score 1 = 7.5uL, 2 = 15ul, 3 = 22uL and 4 = 30uL (1 = 25%, 2 = 50%, 3 = 75%, 4 = 100%). Pool into 1.5mL low bind Eppendorf tubes.*

### AMPure XP clean-up of library pools

*This step removes any primer dimers or unwanted PCR products, by adding 1.0x AMPure XP we are removing anything below 200bp as these fragments are too short to sequence anyway.*

48. Measure the volume in each pool using a pipette (suck up liquid then wind down pipette until liquid is at the bottom then read the volume)
49. Add 1.0x the volume of AMPure XP (room temperature) to each tube, Pipette mix gently 5-10 times
50. Incubate the mixture for 5 minutes at room temperature, flash spin and place on magnetic stand and stand for approximately 5min to capture the bead
51. Aspirate liquid from tubes and discard
52. Wash beads on magnet with 500µL of fresh 70% ethanol
53. Incubate 30 seconds and remove the ethanol
54. Repeat step 52 and 53
55. Remove all residual traces of ethanol
56. Take tube off the magnet and air-dry the beads at room temperature for 3-5 min or until the residual ethanol has completely evaporated. Do not over-dry the beads as

this will decrease yield. The bead pellet is dry when the appearance of the surface changes from shiny to matt

57. Add 35µL 0.1x TE buffer pipette mix to resuspend beads and let stand for 5 min
58. Place tubes on magnet stand for 5 min
59. Carefully pipette supernatant to clean labelled 1.5mL Eppendorf tube - get as much of the buffer as possible. Avoid transferring beads!
60. Library is ready for Hybrid Capture
61. Quantify libraries if necessary

**Hybrid Capture with myBaits® RNA Hybrid Capture probes (Arbor Biosciences)**

*As mentioned earlier in the text, the Hybrid Capture reaction will be performed on pools of libraries to keep costs low. The number of samples in a pool will depend on nature of the project. For ancient DNA studies pools with fewer samples might be more suitable.*

*The following steps are based on the Hybridization Capture for Targeted NGS Manual Version 4.01*
*Because we are using custom made adapters with internal barcodes, we need to use blocker oligos that do match the sequence of our adapters.*

For the Hybrid Capture we follow the manual provided with the baits to the letter. Below, are listed some of the conditions we used for this approach. If the pooling is done as suggested in this protocol- pools of 16 samples- 6 capture reactions will be performed for each bait set. Only 7µL of library is needed per Capture reaction.

- Hybridization was done at 65°C for 48 hours
- Reactions were done in 0.2 mL vessels (PCR strip caps) compatible with magnet stand
- All pipetting was done with multichannel pipettes where possible, using filter-tips
- 20µL of Chill-out™ red liquid wax was added on top of the hybridization mix to avoid evaporation. Tubes were spun shortly to get all liquid below the wax (not part of myBaits protocol!)
- After the 48H the hybridization mix was transferred to a new tube. This was done by pipetting the clear (not red) content into a 200uL pipette filter-tip. It is OK to suck up some of the red wax. At mid-air the library with wax was scrolled up higher into the tip till the wax was all floating on top. Next clear liquid (hybridisation mix) was dispensed in the clean tube by scrolling down the dial of the pipette and stopping just before the wax would hit the tip of the tip

PCR master mix for 6 reactions is needed per Index R1 - Index R2 combination (number of combinations depends on how many unique stubby Y-adapters you have). One for each 16-sample pool. If you are using multiple bait sets with different targets you can use the same Index R1 - Index R2 combinations.

- Library amplification post capture was done with a KAPA HiFi Hot Start Ready Mix PCR Kit. Streptavidin beads were not removed from library

- Post capture indexed primers as listed in Supplemental Table S6 were used for this final PCR stage of the library preparation. All samples should have a unique combination of an internal barcode with Index read1 and read2. Within our lab a barcode-Index 1-Index 2 combination will never be used more than once. All primers have phosphorothioate bonds to protect primers from nuclease degradation
- High copy libraries such as OzBaits_CP libraries generally need 1 or 2 less cycles compared to the Nuclear OzBaits_NR libraries

*Table S3.5 Master mix for final PCR after Hybrid Capture. Index read 1 and read 2 combinations combined with internal 8bp barcode were only used one time ever.*

| ***Cap colour*** | **Reagent** | **Volumes MM 1X** | **X Master Mix +1** |
|---|---|---|---|
| | Enriched library (on streptavidin beads) | 15µL library | |
| | 2X KAPA HiFi HotStart ReadyMix | 25 µL | |
| | Index read 1 primer (P7_primer_Idx###, @ 10 µM) | 2.5 µL | |
| | Index read 2 primer (P5_primer_Idx###, @ 10 µM) | 2.5 µL | |
| | H2O | 5 µL | |
| ***Total*** | | 50 µL | **Dispense 35 µl** |

*Table 3.6 Cycling conditions for post capture PCR using Kapa Hifi Hot Start Ready Mix.*

| | | |
|---|---|---|
| **98 °C** | 2 min | 1 |
| **98 °C** | 20 sec | 14-17 Cycles |
| **60 °C** | 30 sec | |
| **72 °C** | 45 sec | |
| **72 °C** | 5 min | 1 |
| **4 °C** | Hold | |

At the end of this procedure, you should have created libraries with a construct identical to Figure S3.1. These libraries will sequence on any of the Illumina platforms.

#### Post Hybrid Capture PCR clean-up, quantification and size selection

Assuming a full plate of DNA was processed, you should have libraries with 2 different index combinations on Index Read 1 and Index Read 2. For each index combination there are 96 samples which have unique internal molecular identifiers (barcodes) on the first 8 read calls of Read 1 and Read 2. It is essential, for the success of a Hybrid Capture library, to sequence paired-end. In our lab we generally sequence 2X150bp and we aim for approximately 1-2 million reads per sample, this will give plenty of data that is on target and usually will yield most of the chloroplast as a by-product. The inserts are in many cases larger than the 282bp (2x150bp minus 8bpR1 barcode and 8bpR2 barcode and 2x A-tail) sequenced and only the extremities of these inserts are sequenced. It is not necessary for the reads to overlap, this is actual an advantage, as it will allow sequencing outside the targeted regions of interest (the genes for which baits were designed). Generally, a couple of hundred bases into the flanking regions of the genes (introns and spacers), also referred to as the splash zone, will be sequenced with sufficient depth. PCR bias is not a huge problem with this approach as clonal reads can be filtered out easily as the starting material (DNA) for this library was randomly fragmented by sonication.

62. Transfer PCR product with residual streptavidin beads to a new 1.5 mL Eppendorf low bind tube
63. Add 1.0x the volume (50uL) of AMPure XP (room temperature) to each tube, Pipette mix gently 5-10 times
64. Incubate the mixture for 5 minutes at room temperature, flash spin and place on magnetic stand and stand for approximately 5min to capture the beads
65. Aspirate liquid from tubes and discard
66. Wash beads on magnet with 500µl of fresh 70% ethanol
67. Incubate 30 seconds and remove the ethanol
68. Repeat step 66 and 67
69. Remove all residual traces of ethanol
70. Take tube off the magnet and air-dry the beads at room temperature for 3-5 min or until the residual ethanol has completely evaporated. Do not over-dry the beads as this will decrease yield. The bead pellet is dry when the appearance of the surface changes from shiny to matte
71. Add 50ul 0.1x TE buffer pipette mix to resuspend beads and let stand for 5 min
72. Place tubes on magnet stand for 5 min
73. Carefully pipette supernatant to clean well-labelled 1.5mL Eppendorf tube - get as much of the buffer as possible. Avoid transferring beads!
74. Run all libraries through an Agilent 2100 Bioanalyzer with a high sensitivity DNA kit (or similar system, e.g., or Perkin Elmer Labchip GX) and calculate molarity between 300-600bp (range depends on study). If samples are too strong dilute with molecular grade water and run failed samples again
75. Pool libraries to an equimolar concentration. Keep Nuclear (OzBaits_NR) and high copy Chloroplast (OzBaits_CP) libraries separate. We generally pool libraries of 96 samples into one tube before size selection
76. Setup Sage Science Pippin Prep cassette (2% agarose (100bp-600bp)) and set size selecting to 350-600. Depending on library molarity multiple size sections on same pool can be done
77. Recover eluate from elution well in a new 1.5 mL Eppendorf low bind tube

78. Run all size selected libraries through an Agilent 2100 Bioanalyzer again with a high sensitivity DNA kit and calculate molarity
79. Make final library by mixing the Nuclear library with Chloroplast library at a 10:1-2 ratio (10-20% Chloroplast library). Make sure molarity of library is high enough for your sequencing facility. For a Illumina NovaSeq S1 run you'll ideally would generate 30 µL at 10.0nM (Talk to sequencing provider if library too low, they can always play with concentrations).
80. If concentration is too low but volume high enough concentrate library by doing another Ampure clean-up and elute in lower volume, or by evaporation library in a SpeedVac
81. Freeze all intermediate steps of library preparation for troubleshooting of additional sequencing. It is very important to label tubes with as much information as possible, as tubes will have very similar names at each stage of the process. We use small zip-lock bags to keep steps separate
82. Some sequencing facilities might require qPCR quantification before submission. We use the KAPA SYBR® FAST qPCR Master Mix (2X) Kit

*You can pool more than one plate worth of libraries in one pool, sample density will depend on sequencing platform. For an Illumina NovaSeq S1 flow cell we would normally run ~4-5 plates worth of samples (output 650-800M reads), just make sure you use different index combinations*
