## Supplemental 4 for "A hybrid capture RNA bait set for resolving genetic and evolutionary relationships in angiosperms from deep phylogeny to intraspecific lineage hybridization"

**Y-shaped stubby adapter design with molecular identifiers for increased complexity in library design**

### Introduction

The number of applications that now require high-throughput DNA sequencing (HTS) grows steadily with every year and the number of samples that can be processed in one reaction too. When multiple samples are pooled in one single library the identification using molecular identifiers is a fundamental part of the procedure. Pooling samples into libraries has made HTS an affordable approach in research areas that focus on subsets of the genome or technologies that traditionally relied on single sample genotyping or sequencing. Areas such as population genetics, DNA barcoding, forensics are gradually transitioning to HTS technologies to generate their data or are applying new HTS techniques to do research that was previously not possible. Currently the most popular and affordable sequencing platforms are made by Illumina®, with sequencers now able to sequence 16 billion reads on a single flow cell (NovaSeq™ S4 FC, Illumina, San Diego USA, Feb 2019). The standard approach for creating sample unique identifiers to an Illumina library is by adding (through PCR or ligation) one or two indexes to the flanking adapters that contain the flow cell and sequencing primer binding sites. These molecular identifiers are read as separate short (5-10 nucleotide (nt)) reads and are not part of the sequencing output. Standard commercial indexing kits generally contain a set of i5 and i7 adapters or primers with 8 nt indexes that generally combine up to 384 unique index combinations. When a new experiment is designed the same combinations would generally be used again.

One of the main risks in laboratories where libraries are generated on a day-to-day basis are contaminations of previously generated libraries. Ideally NGS libraries are generated without PCR to avoid any PCR bias and basically a fragmented DNA template is ligated with the sequencing adaptors. This is only possible if high quantities of template DNA are present, which is rarely the case. Most library preparations start off at very low molarities and PCR is essential to get libraries to a detectable and measurable concentration. Typical protocols would require multiple PCR steps to finalise a library, so the construct has all the necessary priming, indexing and grafting sites. PCR generates millions and millions of amplicons and these are inevitably dispersed though laboratories as surface contaminations or aerosols through waste (tips, tubes, clean-up wastage), spills or the simple act of opening and closing tubes. If the DNA quality and/or quantity of the tested material is suboptimal this type of contamination might have a huge impact on the final sequencing output. This is basically unavoidable, particularly in a laboratory where many libraries are prepared, and where, usually, related taxa are studied. It is very hard to detect and remove these contaminations when the same index combinations are used over and over again. So, ideally unique index combinations should be used for each individual library, but this is not achievable with the current commercially available two index combinations.

For our laboratory we constructed a library design that has three Molecular Identifiers (MID tags), allowing for 150 million unique combinations, based on 531 8nt MID-sequences with a minimum three base edit distance, as designed by Faircloth and Glenn (1). This document provides a set of robust MID sequences that were tested for power of recovery and avoidance of index read mistakes due to inserts, deletions and substitutions which can happen at each stage of the adapter and library preparation.

The generation of these libraries happens in two stages. First barcoded (8nt) truncated Y-adapters with a thymidine (necessary for ligation to an adenosine overhang) are ligated to fragmented, repaired DNA and amplified with adapter-targeted primers to an intermediate stage. Barcoded samples are then pooled equimolarly for cost-saving purposes and hybrid capture is performed. The Y-adapter designed is shorter (stubby) than the original TruSeq Illumina adapter, this to avoid long indexed adapters, which may interfere with the hybrid capture reaction [2]. In the second stage of the library preparation, in this case the enriched, half-finished products are completed by fusing on the remainder P5 and P7 grafting sequences and indexes with adapter-targeted indexed primers (Supplemental Table S6).
The design of the adapters is inspired on a combination of the adapters used for double digest RAD sequencing [3] and the standard TruSeq™ dual indexing Y-adapters used for Illumina library preparation. The combination of indexed primers and the barcodes allows for pooling of many samples in one sequencing run, reducing the per sample sequencing cost considerably.
Tag or index switching or index hopping, has been a source of concern in recent literature, this is particularly relevant when sequencing amplicons and using a saturated index design [4]. Although not such an issue with the library preparation for shot-gun sequencing library preparation (the basis for a hybrid capture library), the approach presented here would significantly reduce this risk as one single Y-adaptor with a unique barcode will ligate to both ends of the DNA inserts. Bioinformatically this comes down to each barcoded sample having the same 8nt barcode on each end, P7 in forward and the P5 in reverse complement read direction. If swapping would occur, bioinformatic assignment of MID tags to samples would filter reads with nonmatching barcode sequences out.

**Adapter design**

The truncated Y-adapter consists of 2 oligonucleotides that are partially complementary (21 bp, Fig S4.1) a list of 531 pairs of complementary oligonucleotides is presented in Supplementary Table S5. For each barcode two oligonucleotides are needed, the sequences differ only on the insert side of the adapter where the 8nt sequence (based on Faircloth and Glenn (1) design) molecular identifier is located (Figure S4.1). The sequence that contains the 3' end of the barcode has an additional T (thymine) that serves as the complementary base that will anneal with the adenine (A) nucleotide that has been added during the end-repair and A-tailing reaction of DNA fragments (inserts). This same base (T) has a phosphorothioate bond to prevent nuclease degradation. The last base of the 5' end of the barcode has been 5'phosphorylated to allow for DNA ligation. The two tails of the adapter that are not complementary are the adapter priming sites for Read 1 (20bp) and Read 2 (21bp) (Fig S4.1). After ligation the library is amplified to a detectable concentration using primers P7_preCap_Long and P5_preCap_Long (Supplemental Table S6). These are quite long primers and have a melting temperature of ~65°C. We have designed shorter primers (not used in our laboratory) that have a lower melting temperature (Supplemental Table S6). This first PCR will increase the library size with 85bp.

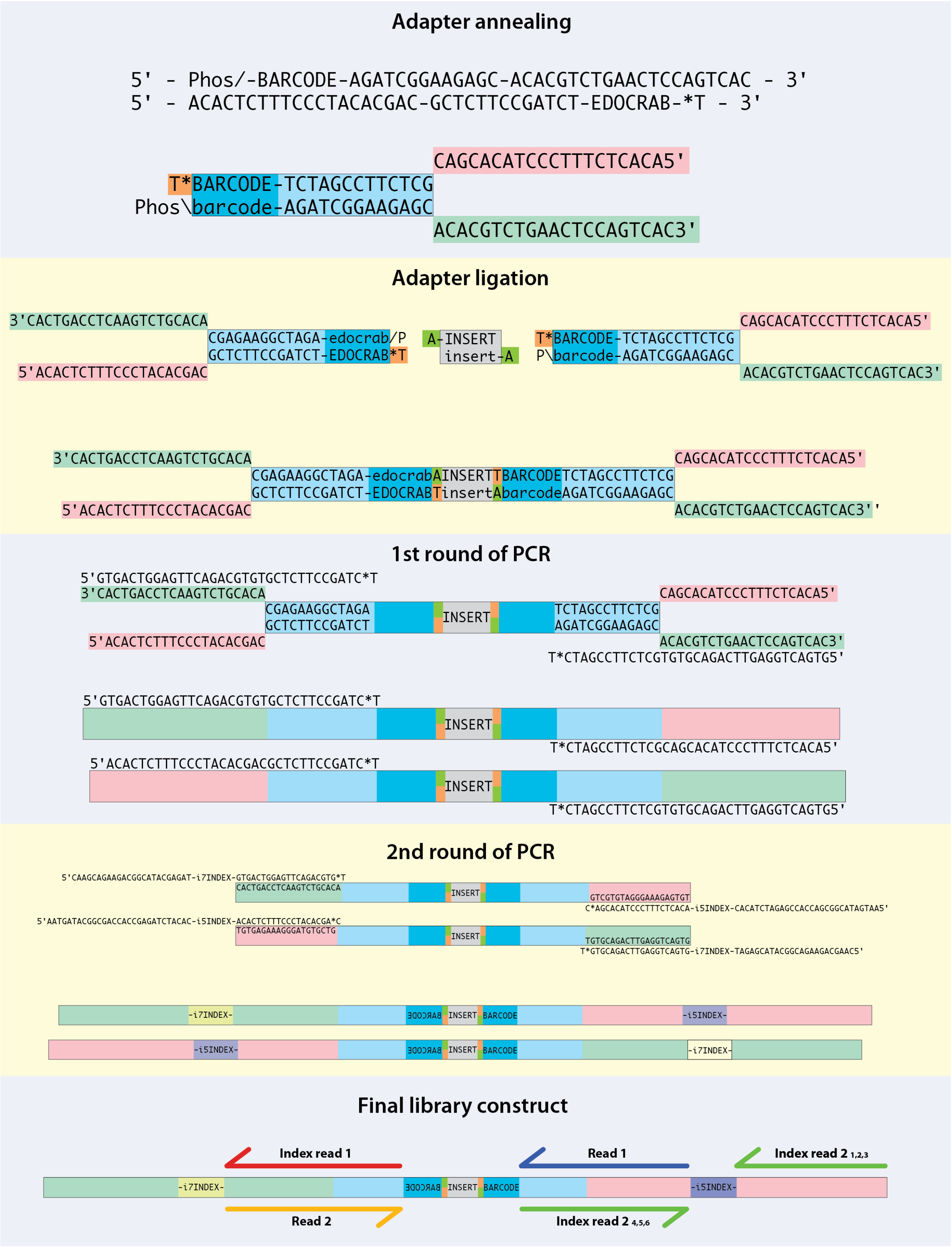


*Figure S4.1 Consecutive stages of stubby Y-adapter construction and library preparation. After adapter ligation and first PCR library is identifiable ready pooling and Hybrid Capture. After enrichment library is finalized by a second PCR, amplifying the P5 and P7 grafting sites on with their respective i5 and i7 indexes. All products will have a P5 and a P7 adaptor on each side of the insert. Phos\=5'phosphorilated base; *= phosphorothioaten DNA bond; Index read 2 $_{1,2,3}$ priming site for MiSeq/HiSeq/NovaSeq; Index read 2 $_{4,5,6}$ priming site for NextSeq, HiSeq 3000/4000*

After the first PCR samples are bioinformatically identifiable and can be pooled to reduce costs of hybrid capture. Depending on how many barcoded stubby Y adapters are available for the study, pooling and second indexed-PCR design will differ. Initially we had 48 barcoded Y-adapters (Tag001-Tag048) available (we now have one plate of 96 Y adaptors) and libraries are generally pooled in lots of 16 samples (6 pools per P5-P7 index combination). The second PCR adds another 69bp to the library. We recommend using 96 barcoded Y-adapters (Tag001-Tag096) as this will make the method easier and more robust.

Faircloth and Glenn (1) provided 531 8nt sequences with a 3 edit distance (see Supplemental Table S7), but this list was provided in alfabetical order. We randomized the adapters to avoid all barcodes staring with the same nucleotides. This is essential as cluster coordinates are determined based on the images from the first four sequencing cycles and lack of sequence diversity might lead to cluster identification problems.
For this library design to work effectively paired-end sequencing is essential. We recommend at least 150 paired-end sequencing, as nine base pairs of each read are lost (8bp of barcode plus T for ligation), leaving 141 bases for read 1 and read 2. For hybrid capture libraries were size selected to 350-600bp, removing the 154bp of non-insert adapter DNA would leave an insert size of approximately 200-450bp.
It is important to keep a thorough record of the P5-Index-P7-Index-Barcode combinations used in a laboratory, if a contamination occurs, events can be tracked down to the original source and eliminated.

**Conclusion**
The library design presented is an affordable modification to standard laboratory practices that allows for much higher complexity in a multiplex NGS experiment design. Laboratories that strive to use unique MID tag combinations for each individual sample run out of options very quickly with the limited number of combinations possible using commercially available indexing kits. This is particularly a problem when NGS platforms can generate higher read numbers with each new development. In fields like forensics and ancient DNA research where trace DNAs might be of significance importance, both positively and negatively. Being able to keep track of these bioinformatically can be of significant use.
There are also obvious down sides of using this approach, particularly if short read chemistry is used, if for example an 8nt MID is used 9 bases of a read would be lost. If running 75 cycle chemistry this would be a loss of 10% of the data.
This approach is particularly useful if a high number of samples is processed, the number of adaptors used in one run should be high to avoid clusters recognition failure.
